# On the overlap between scientific and societal taxonomic attention - insights for conservation

**DOI:** 10.1101/334573

**Authors:** Ivan Jarić, Ricardo A. Correia, David L. Roberts, Jörn Gessner, Yves Meinard, Franck Courchamp

## Abstract

Attention directed at different species by society and science is particularly relevant within the field of conservation, as societal preferences will strongly impact support for conservation initiatives and their success. Here, we assess the association between societal and research interests in four charismatic and threatened species groups, derived from a range of different online sources and social media platforms as well as scientific publications. We found a high level of concordance between scientific and societal taxonomic attention, which was consistent among assessed species groups and media sources. Results indicate that research is apparently not as disconnected from the interests of society as it is often reproached, and that societal support for current research objectives should be adequate. While the high degree of similarity between scientific and societal interest is both striking and satisfying, the dissimilarities are also interesting, as new scientific findings may constitute a constant source of novel interest for the society.

## I. INTRODUCTION

Species receive uneven attention in terms of scientific research (Clark and May 2002; Proenca et al. 2008; De Lima et al. 2011; Murray et al. 2015; Donaldson et al. 2016; Fleming and Bateman 2016). This uneven scientific focus is driven by diverse factors, such as geographic location, species accessibility, suitability for use as model species, conservation status, and researchers’ own personal interests (Jarić et al. 2015). Society, however, can also influence research focus through policy and funding agendas, while science in turn influences societal attention through scientific communication and media representation. Contrastingly, choices of studied species are sometimes criticized as leading to a waste of societal resources when they do not appear to match the immediate interest of taxpayers.

Based on the main drivers of societal and scientific taxonomic attention identified so far in the literature, we suggest that there are at least three general categories of drivers of societal and scientific taxonomic attention: 1) intrinsic, species-related factors, which can also be considered as elements of species charisma, 2) population-level or spatial factors, and 3) socio-economic factors. Major intrinsic factors include body size, unique morphology, distinctive coloration patterns, anthropomorphism, behavior, social structure and neotenic features (Moustakas and Karakassis 2005; Stokes 2007; Wilson et al. 2007; Martín-FÓres et al. 2013; Żmihorski et al. 2013; Kim et al. 2014). Other recognized proxies for scientific and societal taxonomic preferences are phylogenetic distance from humans and structural complexity (Proenca et al. 2008; Martín-LÓpez et al. 2011; Martín-FÓres et al. 2013), although both are associated with already listed factors such as anthropomorphism and body size. Population-level or spatial factors include abundance, range size, range proximity to or overlap with developed nations, extinction risk, and habitat accessibility (Wilson et al. 2007; Brooks et al. 2008; Sitas et al. 2009; Trimble and van Aarde 2010; Fisher et al. 2011; Żmihorski et al. 2013; Dos Santos et al. 2015; Jarić et al. 2015; Zhang et al. 2015). Socio-economic factors are represented by the species economic value (e.g. as an object of trade or tourism), its pest status, potential threat to humans (e.g. venomous or aggressive species), presence of key ecological values or ecosystem services, and various cultural values (i.e. traditional, religious, etc.) (Moustakas and Karakassis 2005; Wilson et al. 2007; Proenca et al. 2008; Jarić et al. 2015; Zhang et al. 2015; Donaldson et al. 2016; Roll et al. 2016).

While previous research has addressed the factors underlying uneven taxonomic attention, the actual level of overlap between societal and scientific attention has been poorly quantified. In the current information age, society has access to and produces much more content than any previous generation. Due to the sheer amount of accessible information, it becomes necessary to make choices regarding the attention scope. Consequently, it may be interesting to compare the species chosen by scientists and by the rest of the society. This question was previously addressed in the seminal work of Wilson et al. (2007), however this was based on a rather limited sample. While it has not received further attention so far, this issue remains highly relevant, particularly within the field of conservation biology. As stated by Stokes (2007), societal preferences are just as important for the success of conservation efforts and survival of many endangered species as are common ecological determinants, such as minimum population size and habitat requirements. Societal preferences can play a wide range of roles. People express their views and interests using various widespread media, and not all have the opportunity to express their interest in a more active way, such as engagement in conservation non-profit organizations. Societal attention towards particular species can be beneficial if it helps society to understand the need for conservation action and to support it. Approaches that aim to attract societal attention towards conservation goals, such as flagship species concept, have proven to be successful in attracting societal support and funding (Veríssimo et al. 2011, 2017). On the other hand, increased attention might sometimes lead people to exert increasing negative pressure on the species they are interested in, akin to the Anthropogenic Allee Effect (Courchamp et al. 2006), or alternatively to contest actions against invasive alien species (Courchamp et al. 2017).

Here we take advantage of emerging culturomic techniques (Michel et al. 2011; Ladle et al. 2016; Sutherland et al. 2018) to assess the similarities and differences in the societal and scientific interests in different species, based on scientific publications and a range of different online sources and social media. We assessed the relationship between the scientific and societal taxonomic attention within four species groups that predominantly consist of charismatic and threatened animals: carnivorans, primates, marine mammals and birds of prey. We discuss the drivers of observed relationships and overlaps, and address their implications for conservation planning and management.

## II. METHODS

In order to account for the problems associated with vernacular species names, such as synonyms, homonyms and multiple meanings, we used the approach proposed by Jarić et al. (2016) and Correia et al. (2017). Species lists, comprising diurnal birds of prey (orders Accipitriformes, Falconiformes and Cathartiformes), Carnivora, Primates and marine mammals (cetaceans and pinnipeds), were obtained from the IUCN Red List database (IUCN 2017). Extinct species and those described after 1995 were excluded from the analysis, which resulted in a total of 1058 species in the dataset (318 birds of prey, 252 carnivorans, 370 primates and 118 marine mammals). Search of scientific publications and online media was conducted by using species scientific names and scientific synonyms, each placed in parentheses, within a same search query (i.e., [“*species name*” OR “*synonym #1*” OR “*synonym #2*” OR…]). This resolved the problem of potential double entries, and the results were thus expressed as the number of unique records per species.

Research attention was defined as the number of scientific articles indexed within the Web of Knowledge (available at www.isiknowledge.com) for a given species. The search was conducted within titles, abstracts, and keywords of referenced publications published during 1996-2016. Keywords that are automatically assigned by the Web of Knowledge (i.e. Keywords Plus) were not considered in the analysis, due to their low reliability (Wilson et al. 2007; Fisher et al. 2011).

Media coverage for each species was estimated based on the following five online sources: Internet pages containing the species name, online articles in selected major international newspapers (The New York Times, The Guardian, Le Monde, Washington Post, and Asahi Shimbun), Twitter, Facebook, and pictures posted on the Internet for each of the studied species (Jarić et al., 2016). Media coverage data collection was performed in line with the approach by Correia et al. (2017), by using the Google’s Custom Search Engine API. Searches were carried out during June 2017, with search queries for each of the online sources based on Jarić et al. (2016): 1) Internet pages – [“*species name*”], 2) Twitter – [“*species name*” site:twitter.com], 3) Facebook – [“*species name*” site:facebook.com], 4) Newspapers – [“*species name*” (site:nytimes.com OR site:theguardian.com OR site:lemonde.fr OR site:washingtonpost.com OR site:asahi.com)], and 5) Photographs – [“*species name*” (filetype:png OR filetype:jpg OR filetype:jpeg OR filetype:bmp OR filetype:gif OR filetype:tif OR filetype:tiff)], where species name was represented by both scientific name and scientific synonyms.

The resulting dataset features the number of records per species and per assessed sources. Since the variables were not normally distributed (Kolmogorov-Smirnov test, *p*<0.001), nonparametric tests were applied. Relationship between the number of scientific publications and the five online media sources, within each of the four studied species groups, was assessed using a Spearman’s Rank test, with Bonferroni correction. We also conducted ranking, by ordering species based on the number of results for each of the five online media sources assessed, and estimating the average rank across the sources; ranking was also performed for scientific publications.

## III. RESULTS

The average number and range of records obtained for each species group, for scientific publications and each of the five assessed online media sources, are presented in Table S1 (Supplementary material). Results indicated strong correlations (0.751 mean correlation coefficient, *p*<0.001) between the number of scientific publications per species and the number of results from each of the online media sources assessed, in each of the four studied species groups (Fig. 1; Table I). Correlations were strongest in carnivorans and lowest in primates (0.836 and 0.696 mean correlation coefficients, respectively). Regarding the media sources assessed, correlations with the number of scientific articles per species were strongest for Internet pages and lowest for newspaper articles (0.889 and 0.550 mean correlation coefficients, respectively; Table I). All correlations remained significant following a Bonferroni correction.

**Fig. 1.**
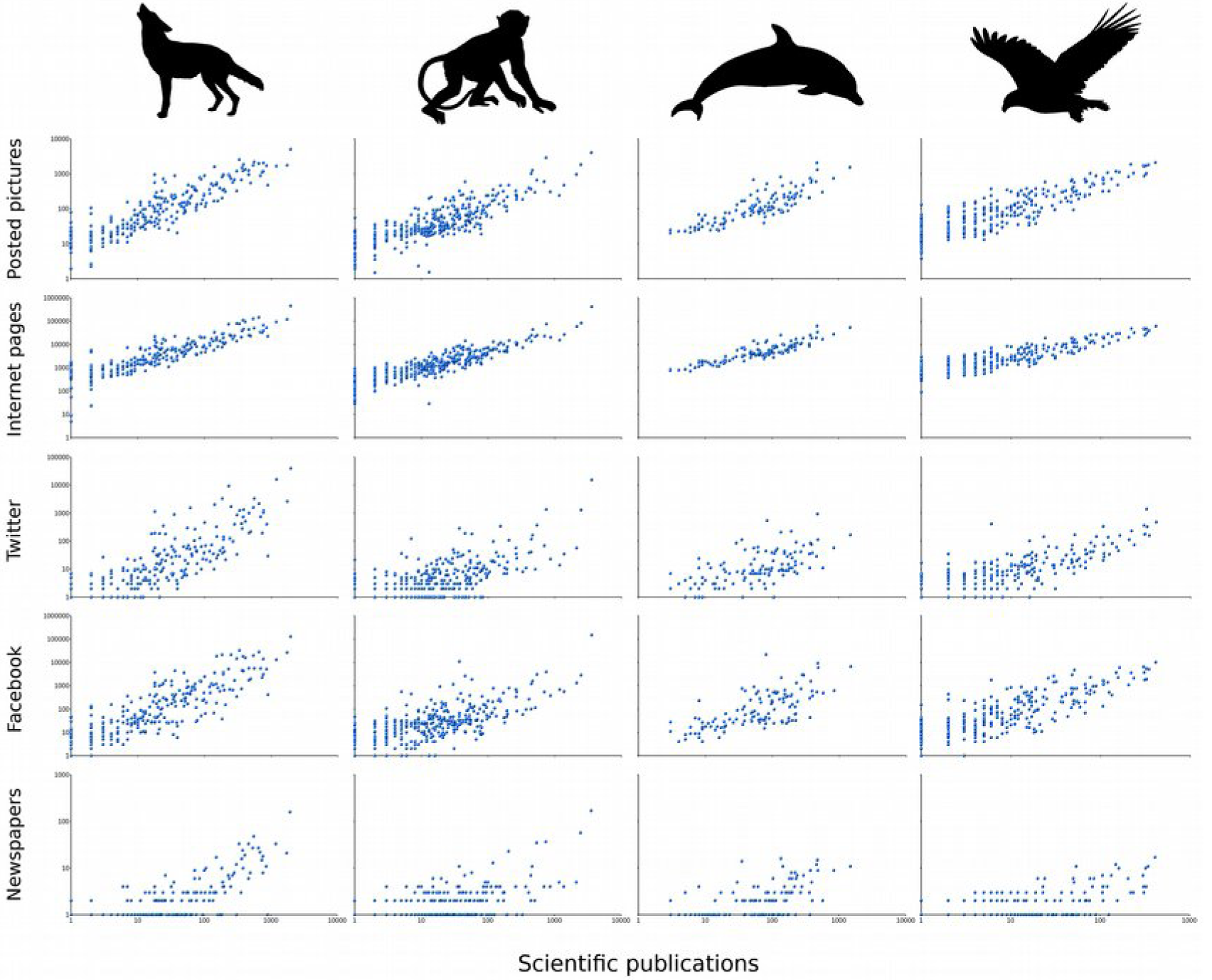
Relationship sdf between the scientific attention (Web of Science) and coverage within different media sources (Internet pages, Twitter, Facebook, newspapers, and photographs posted on the internet) in different species groups (Carnivora, Primates, marine mammals and birds of prey); axes represent logarithmic scales. Presented data were transformed using x ← x + 1, in order to allow presentation in log-plots of results with the value of zero.

**TABLE 1.**
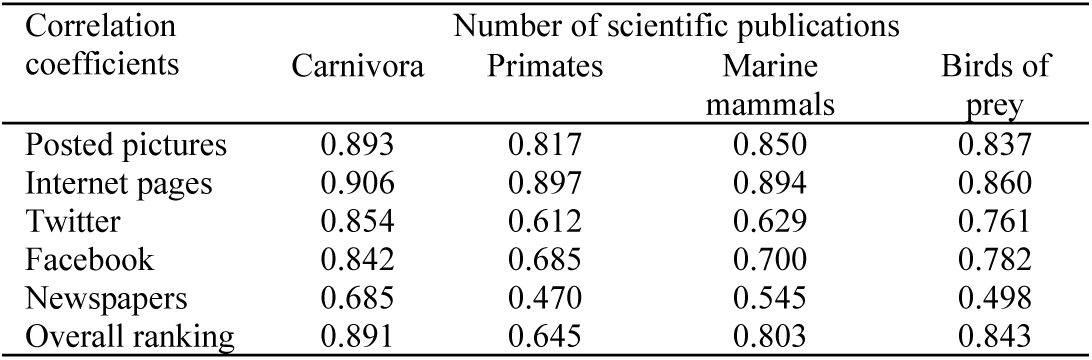
RELATIONSHIP BETWEEN THE SCIENTIFIC ATTENTION (WEB OF SCIENCE) AND COVERAGE WITHIN DIFFERENT MEDIA SOURCES (INTERNET PAGES, TWITTER, FACEBOOK, NEWSPAPERS, AND PHOTOGRAPHS POSTED ON THE INTERNET) IN DIFFERENT SPECIES GROUPS (SPEARMAN’S NON-PARAMETRIC CORRELATION TEST, *P*<0.001; ALSO SEE FIG. 1); SEE THE TEXT FOR INFORMATION ON OVERALL RANKING APPROACH.

| Correlation coefficients | Number of scientific publications |  |  |  |
| --- | --- | --- | --- | --- |
|  | Carnivora | Primates | Marine mammals | Birds of prey |
| Posted pictures | 0.893 | 0.817 | 0.850 | 0.837 |
| Internet pages | 0.906 | 0.897 | 0.894 | 0.860 |
| Twitter | 0.854 | 0.612 | 0.629 | 0.761 |
| Facebook | 0.842 | 0.685 | 0.700 | 0.782 |
| Newspapers | 0.685 | 0.470 | 0.545 | 0.498 |
| Overall ranking | 0.891 | 0.645 | 0.803 | 0.843 |

Overall species ranks within social media had strong positive correlations with their ranking based on scientific publications (Table I). Lists of top-ranked species based on their overall presence in social media were fairly similar to those that reached top ranks within scientific publications (Table II). Common bottlenose dolphin (*Tursiops truncatus*) was the most popular marine mammal species within the scientific community, and the second-highest ranking marine mammal species for the general society. Top-ranked birds of prey in science and among the general society, as well as top-ranked carnivorans in science, are exclusively represented by European and North American species. On the other hand, top-ranked carnivorans among the general society also comprised two big cats from Africa and Asia, lion (*Panthera leo*) and tiger (*P. tigris*). Top-ranked primates were dominated by macaque species (*Macaca sp.*) such as rhesus macaque (*Macaca mulatta*), the highest ranked primate within both sources, as well as by big apes (Table II).

**TABLE II.**
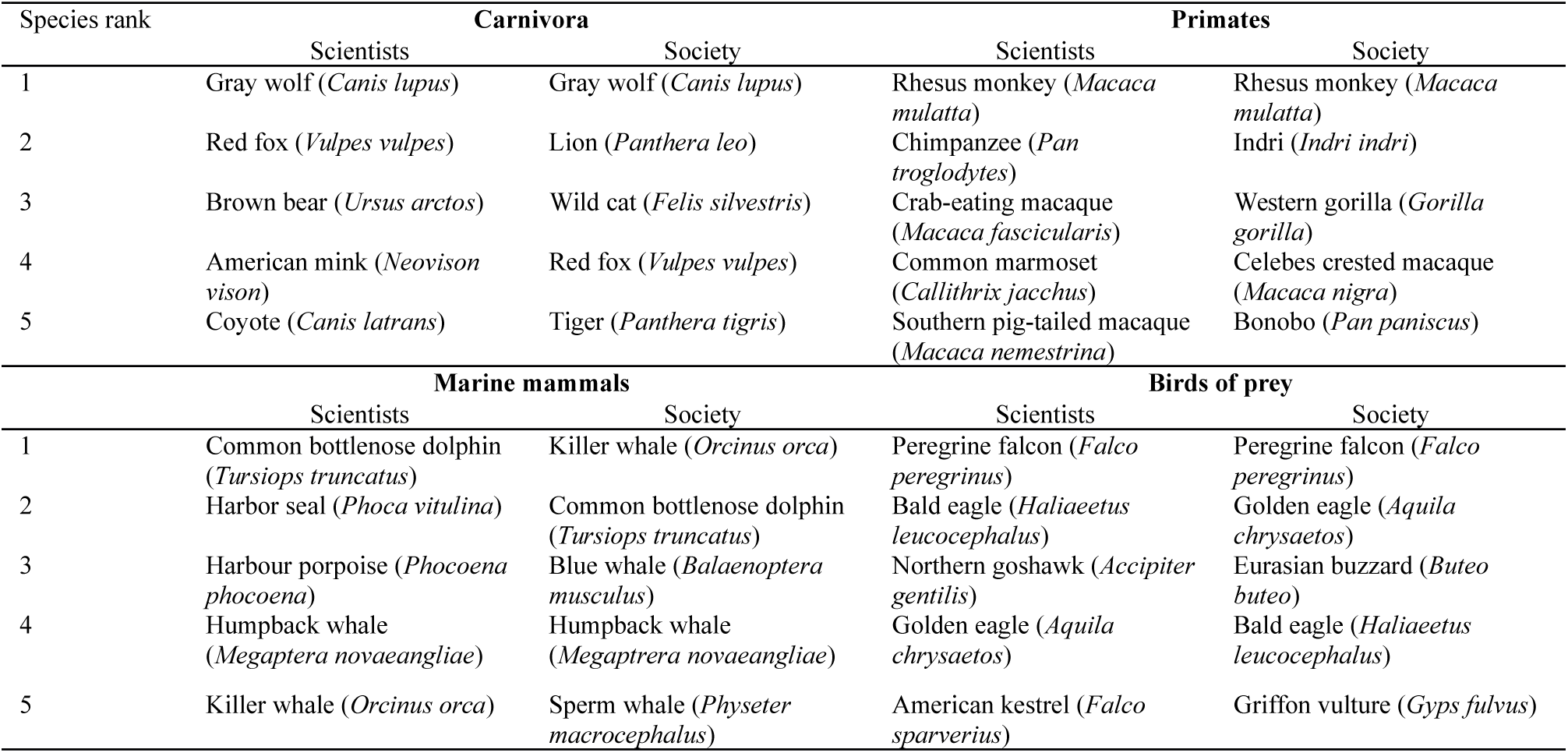
TOP FIVE RANKED SPECIES FROM THE FOUR ANALYZED SPECIES GROUPS, BASED ON THE FREQUENCY OF THEIR PRESENCE IN SCIENTIFIC PUBLICATIONS AND THE LEVEL OF SOCIETAL ATTENTION, ESTIMATED AS THE AVERAGE RANKING ACROSS FIVE ASSESSED ONLINE SOURCES.

## IV. DISCUSSION

The literature indicates that species coverage may differ among different media (Jacobson et al. 2012). However, in our study all five assessed online media sources provided similar results, which suggests that they can potentially be used interchangeably as a measure of societal taxonomic attention. Yet, most of them either represent specific sectors, such as newspaper articles, or are generated by different processes, and therefore may provide essentially different information. Although they have been relatively rarely used so far, web-based images also seem to represent a suitable tool for data mining (Barve 2014; Jarić et al. 2016; Ladle et al. 2017; Sherren et al. 2017a, 2017b). Images and other visual media may be especially adequate for culturomic studies that are focused on species attractiveness and charisma, which is particularly relevant for the field of conservation biology. As our study demonstrates, the use of images within this field can go beyond the analysis of cultural ecosystem services (Willemen et al. 2015; Martínez Pastur et al. 2016; Hausmann et al. 2017). Use of social media in conservation science is still somewhat limited (Di Minin et al. 2015), but is rapidly increasing. Twitter and Facebook represent dominant social media platforms, which makes them suitable research tools (Miller 2011; Roberge 2014; Papworth et al. 2015). They are rapidly changing communication and information sharing dynamics, and are increasingly used as communication platforms by the scientific community and other groups from the biodiversity conservation field (Naaman et al. 2011; Bombaci et al. 2015). Online news media have a wider reach than traditional printed newspapers, and are considered suitable to reflect societal attention and popular attitudes (Veríssimo et al. 2014; Papworth et al. 2015). However, we observed very low presence of species in newspaper articles (Table S1). As much as 68% of the assessed species had no newspaper articles, while only 20% of the species had more than a single result. This issue may be partly due to the search conducted by using only scientific species names, although such approach has been validated (Jarić et al. 2016) and does not seem to be an issue with the other online sources, such as web-based images. To a certain extent, this may be due to news media commonly focusing on only a small proportion of high-profile species (i.e., charismatic species, or those with high economic value), while the majority of other species seem to end up being neglected. Additionally, low species coverage by online news media may stem from inappropriate publishing practices. Wildlife observers or photographers often strive to provide the scientific name of the species they are posting about on the web, while journalists do not. Due to potential implications for science education and societal outreach, it would be valuable to explore this issue further.

Based on all these various representations of societal attention, our analysis unveiled a high level of concordance between scientific and societal taxonomic attention, and this was consistent among assessed species groups and online media sources. This shows that scientific focus is not remote from societal attention towards different species, and *vice versa*, a finding also reported by Wilson et al. (2007). On the one hand, this can be interpreted as a positive outcome, since scientists are apparently well aligned with societal attention, which is what the general society, as providers of public funding, and consequently the funding agencies, would request. On the other hand, if research focus and societal attention are both considered to be biased (Clark and May 2002; Sitas et al. 2009; Kim et al. 2014; Roberge 2014; Donaldson et al. 2016; Wilson et al. 2016; Troudet et al. 2017), it is of special importance to understand the mechanisms that produce such biases. They are likely represented by a similar set of drivers that are influencing societal and scientific attention, as well as by the interaction between the two groups. However, as stated by Troudet et al. (2017), while the presence of interaction between the scientists and the general society is not questionable, it remains particularly challenging to clarify the actual direction and causality of influence between the two groups. It is important to emphasize that our study focused only on the level of overlap among the coverage of different media sources and scientific publications, and not on the actual media content or mechanisms that are driving public and scientific attention.

The top five ranked species in each of the four studied species groups revealed a substantial overlap between scientific and societal focus (Table II). Popularity of common bottlenose dolphin within the scientific community is mainly due to its use as a model species in experiments on a wide range of topics, such as echolocation, behaviour, intelligence, swimming and communication (Jarić et al. 2015). At the same time, its popularity for the general society probably comes from its ubiquity and high presence in both captivity and popular culture. Dominance of European and North American birds of prey and carnivores points to commonness and range overlap with developed countries as major drivers of taxonomic attention. Lion and tiger, African and Asian carnivores that were among the top-ranked species by the general society, were previously identified as the two most charismatic animals globally, while the gray wolf (*Canis lupus*), highest-ranked carnivoran within both sources in the present study, was identified in the same study as the 9^th^ most charismatic animal (Courchamp et al., unpublished). Among the primates, those that are prominently used as model species dominated the ranking. Based on the individual checking of internet sources and online news for the rhesus macaque, it seems that its high presence in online media is mainly due to health- and medicine-related content, where the species is mentioned as a study system (e.g. efforts at developing HIV vaccine). Big apes are among the most charismatic primate species (Courchamp et al., unpublished), mainly due to higher levels of anthropomorphism and comparatively larger body size than in other primates. Use as model species is likely a less important attention driver for the society, which may explain why big apes are more prominent among the top-ranked primates for the general society than for science. This might have also contributed to the weaker correlations between the societal and scientific attention for primates than in the other three assessed species groups.

It is important to note the potential risk of a statistical bias when using a measure of societal interest that depends on the capacity of people to interact with the Internet. Users from developed countries and the related content are likely to be overrepresented, and those regions are also where most of the scientific output originates, which might make species from those areas also more prominent in both of the assessed sources (Martin et al. 2012; Amano and Sutherland 2013; Amano et al. 2016; Wilson et al. 2016). This calls for caution when interpreting presented results. On the other hand, the potential problem of biased media coverage when focusing studies of online media on only a single language (Bhatia et al. 2013; Funk and Rusowsky 2014) was resolved here by using scientific names as search keywords (Jaric et al. 2016; Correia et al. 2017).

Understanding of societal and scientific attention is especially relevant within the field of biodiversity conservation, due to its potential impact on the general support for conservation efforts (Stokes 2007). The biodiversity conservation arena is generally considered to be represented by four distinct, interacting sectors: the scientific community, policy makers, news media, and the general society (Papworth et al. 2015). The extent to which the four sectors align in their focus depends on their sensitivity and susceptibility to each of the three general taxonomic attention drivers listed in the introduction (i.e., intrinsic, population-level/spatial, and socio-economic), as well as on the level of inter-sectoral interaction. The scientific community is strongly influenced by research funding and science policy. If both funding and the policy follow wider societal preferences, such as species charisma, scientific attention will correspond well to that of the general society. Scientists in turn also influence societal interests by communicating information and new knowledge to the general society, both directly, through different outreach activities, and indirectly, through news media and by informing and guiding policy development and conservation decision making (Moustakas and Karakassis 2005; Trimble and van Aarde 2010; Papworth et al. 2015). Certain levels of dissimilarity between scientific and societal attention can also produce positive effects, by bringing new centers of interest to the general society. Each of the sectors is also subject to its own internal mechanisms that generate or maintain existing taxonomic attention patterns. For example, research inertia may contribute to perpetuated biases in taxonomic attention in science (Jarić et al. 2015; Troudet et al. 2017). Researchers often focus on well-studied species they are familiar with, with proven potential to attract funding, and past research in one area will therefore have a tendency to generate more research in the same area (Martín-LÓpez et al. 2009; Dos Santos et al. 2015; Correia et al. 2016a). Biased scientific publishing practices, such as “taxonomic chauvinism” (Bonnet et al. 2002), will also contribute to maintaining taxonomic biases in research.

The same drivers of taxonomic attention can impact both scientific community and the general society, while working within each of the two sectors through different mechanisms. For example, for many species, range proximity and population abundance seem to be two important drivers of societal attention, recognized as species commonness or familiarity (Żmihorski et al. 2013; Schuetz et al. 2015; Correia et al. 2016b). At the same time, they are also relevant drivers of scientific attention, by contributing to improved species accessibility, reduced logistical challenges and lower research costs (Dos Santos et al. 2015; Jarić et al. 2015). It is also important to bear in mind that scientists also represent members of the general society, with their own interests and susceptibility to drivers such as species charisma (Lawler et al. 2006; Lorimer 2007; Wilson et al. 2007; Smith et al. 2009). It is therefore possible that, in cases where liberty of choosing research topic exists, societal and scientific interests will essentially be the same.

One implication arising from the results is that environmental education projects or programs should target species beyond the focus of the general society, to allow the discovery and promote interest in such species. Another alternative would be to focus on the very species that both scientist and the general public are interested in, provide more knowledge on those species, and thus further strengthen societal support for current research efforts. One of the often-advocated measures in this respect is to intensify and improve the effectiveness of science communication (Dietz 2013). However, for science communication to accomplish the desired aims, a first step would be to consider mechanisms that shape societal attention, and to ensure that science outreach initiatives are structured based on identified societal beliefs, values, information gaps and misconceptions (de Bruin and Bostrom 2013). Meanwhile, and despite its biases, the scientific community and conservationists should try to make the most of existing societal attention by taking advantage of flagship species to attract conservation funding and support for a wider range of species (Clark and May 2002; Jepson and Barua 2015).

## V. CONCLUSIONS

Societal interest in the fate of endangered species is a crucial prerequisite for effective conservation programs, given that the general society is likely to protect only what it recognizes as important (Stokes 2007; Kim et al. 2014). Societal awareness and societal values will largely determine whether conservation initiatives will receive necessary support and lead to adequate policy change (Papworth et al. 2015). On one hand, societal attention is closely associated with scientific attention, which should ensure that the societal support for current research objectives should not be lacking. This also implies that scientists are not so disconnected from the rest of society. On the other hand, societal and scientific interests are not perfectly aligned, which indicates that there is room for studies of species not *a priori* interesting to the society. In fact, scientists may still remain free of the potential biases of societal taxonomic interests, while they are at the same time in good position to provide novel knowledge and new points of interest to the society.

## ACKNOWLEDGMENTS

IJ acknowledges the sponsorship provided by the The J. E. Purkyně Fellowship of the Academy of Sciences of the Czech Republic, Alexander von Humboldt Foundation and the German Federal Ministry of Education and Research (BMBF), as well as by the Project No. 173045, funded by the Ministry of Education, Science and Technological Development of the Republic of Serbia. RAC is currently supported by a post-doctoral grant from Fundação para a Ciência e Tecnologia (SFRH/BPD/118635/2016). FC acknowledges the support by the Invacost research program (ANR and BNP Paribas).

